# Pre-existing stem cell heterogeneity dictates clonal responses to acquisition of cancer driver mutations

**DOI:** 10.1101/2024.05.14.594084

**Authors:** Indranil Singh, Daniel Fernandez Perez, Pedro Sánchez, Alejo E. Rodriguez-Fraticelli

## Abstract

Cancer cells display wide phenotypic variation even across patients with the same mutations. Differences in the cell of origin provide a potential explanation, but these assays have traditionally relied on surface markers, lacking the clonal resolution to distinguish heterogeneous subsets of stem and progenitor cells. To address this challenge, we developed STRACK, an unbiased framework to longitudinally trace clonal gene expression and expansion dynamics before and after acquisition of cancer mutations. We studied two different leukemia driver mutations, Dnmt3a-R882H and Npm1cA, and found that the response to both mutations was highly variable across different stem cell states. Specifically, a subset of differentiation-biased stem cells, which normally become outcompeted with time, can efficiently expand with both mutations. Npm1c mutations surprisingly reversed the intrinsic bias of the clone-of-origin, with stem-biased clones giving rise to more mature malignant states. We propose a clonal “reaction norm”, in which pre-existing clonal states dictate different cancer phenotypic potential.

**Highlights:**

- Single cell tracing of cancer initiation at the clonal level (STRACK).
- Ex vivo expansion cultures sustain intrinsic and heritable HSC heterogeneity.
- Premalignant mutations enhance the self-renewal of high-output stem cells, increasing their survival probability.
- Transforming mutations reprogram low-output stem cell fates to more mature malignant states.

## Introduction

Cancer cells display striking phenotypic variation, within and across patients, yet the origins of this variation are still unclear (Lenz et al. 2022). Since cancer is a clonal disorder, arising from a single cell, researchers have long hypothesized that phenotypic heterogeneity could be a consequence of the cell type that acquires the driver mutations (Blanpain 2013; Visvader 2011). The “cell-of-origin” model transformed the cancer field, leading to transformational discoveries across various tumors and cell types (Baggiolini et al. 2021; Rajbhandari et al. 2023). However, these classic cell-of-origin studies have several limitations. Firstly, they usually induce mutations at a population level, lacking information about individual cell heterogeneity. Second, they rely on reporter genes or surface-markers to isolate the cell population of interest, and are thus biased by the tools and prior knowledge of the model system. Finally, there are currently no methods that capture, with high resolution, the pre-existing states (and fates) of the single cells that give rise to the malignancy after mutagenesis. Solving these limitations could be critical to deepen our understanding of cancer initiation mechanisms.

The cell-of-origin hypothesis has been extensively characterized in cancer types with few driver mutations, such as myeloid malignancies. Depending on whether mutations are introduced in the hematopoietic stem cells (HSCs), at the top of the hematopoietic hierarchy, or in the more mature myeloid progenitors (MPs), researchers have consistently shown differences in the resulting phenotypes (A. V. Krivtsov et al. 2013; SanMiguel, Eudy, Loberg, Miles, et al. 2022; Stavropoulou et al. 2016; Cai et al. 2020; Zeisig et al. 2021; Taussig et al. 2010; Cozzio et al. 2003; Huntly et al. 2004; Andrei V. Krivtsov et al. 2006; George et al. 2016). However, recent single-cell sequencing studies have elucidated that stem and progenitor cell populations are highly heterogeneous and cannot be simply dissected through surface markers (Paul et al. 2016; Giladi et al. 2018). Furthermore, we and others have shown that even the HSCs, at the top of the hierarchy, are long-term biased at the level of both state and function, with a multiplicity of fate-imprinted clonal hierarchies likely co-existing in the bone marrow (Rodriguez-Fraticelli et al. 2020; L. Li et al. 2023; Meng et al. 2023; Jindal et al. 2023; Perié et al. 2015; Wang et al. 2022; Wagner and Klein 2020; Weinreb et al. 2020; Rodriguez-Fraticelli et al. 2018; Tian et al. 2021; Naik et al. 2013; Dykstra et al. 2007; Wilson et al. 2015). Yet, due to the lack of high-resolution cell-of-origin techniques, the functional significance of stem cell heterogeneity in tumor initiation remains poorly understood (Haas, Trumpp, and Milsom 2018), leaving researchers to rely solely on inference (Tong et al. 2021).

Here, we present a system called STRACK (Simultaneous Tracking of Recombinase Activation and Clonal Kinetics), that precisely addresses these knowledge gaps, unbiasedly linking pre-existing stem cell states (and intrinsic fates) with their potential cancer states and fates. STRACK takes advantage of defined primary stem cell culture systems to explicitly minimize the confounding effect of extrinsic variables and solely focus on intrinsic determinants. To this end, STRACK combines long-term *ex vivo* PVA-based expansion cultures, which can sustain and expand HSCs and myeloid progeny for weeks (Wilkinson et al. 2019; Che et al. 2022), mouse models carrying different Cre/Flp-inducible leukemia mutations (Loberg et al. 2019), a new palette of LARRY expressed barcode libraries to track clones (Weinreb et al. 2020), and a sister-cell clone splitting strategy (Weinreb et al. 2020; Tian et al. 2021). This unique combination of methods allowed us to sample the system longitudinally and obtain a dense clonal and transcriptional landscape for the same set of clones, with and without mutations.

## Results

### PVA-based expansion cultures maintain heterogeneous and deterministic clonal behaviors

In order to characterize HSC clonal behaviors in PVA-based *ex vivo* expansion cultures, we profiled thousands of HSC clones through a 27-day protocol using single-cell lineage tracing and RNA profiling. For this, we genetically labeled ∼10,000 long-term hematopoietic stem cells (Lineage- Sca-1+ cKit+ CD48- CD150+ CD201+ or “E-SLAM”) with new LARRY barcoding libraries expressing the mT-Sapphire fluorescent protein (Figure S1A). Labeled HSCs were expanded across multiple wells for 27 days, sorted and randomly sampled for scRNAseq as depicted in the schematic at day 7, 14 and 27 (Figure 1A and Supplementary Table 1 ∼see methods)(Weinreb et al. 2020).

**Figure 1.**
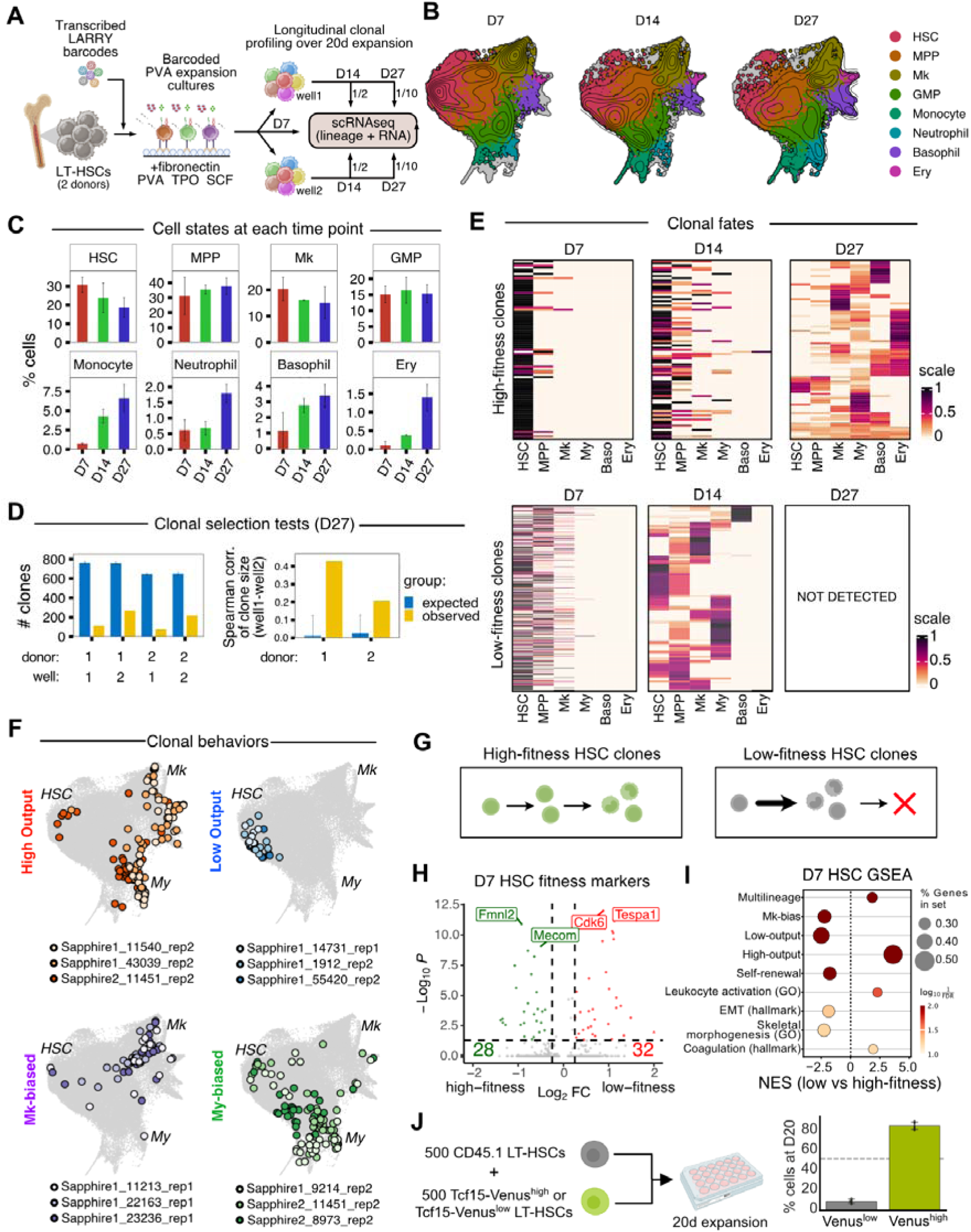
State-fate analysis in *ex vivo* HSC expansion cultures. A) Experimental design for state-fate analysis in *ex vivo* expansion cultures. B) UMAP of integrated data from day 7, day 14 and day 27 sampling timepoints. C) Distribution of cell state proportions across cluster groups in each sampling timepoint. D) Observed versus expected number of clones based on a stochastic sampling model. Right, spearman correlation of sister-cell clone sizes across split independent wells (observed, yellow; blue, expected). E) State-bias heatmaps showing clones (rows) and clusters/states (columns), colored by the intra-clonal fraction in each cluster/state. Clones shown are all those detected in both day 7 and day 14. Clones are separated into two heatmaps (top - bottom), depending on whether the clone is detected (in at least 2 cells) at day 27. F) UMAP showing example clonal behaviors at day 27 from different experiments/replicates. G) Scheme of the interpretation of results based on clonal groupings. H) Volcano plot for day 7 HSCs comparing low-vs-high fitness clonal groups. Selected genes are highlighted. Markers associated with each group are shown in green (high-fitness) or red (low-fitness). I) GSEA of hallmarks, GO-terms and HSC signatures for the differential gene expression results of high versus low-fitness HSCs at day 7. J) Validation experiment using competitive *ex vivo* HSC expansion of Tcf15-Venus^high^ versus Tcf15-Venus^low^ cells.

Most of the cells profiled on day 7 expressed markers of HSCs (*Procr, Hlf, Mecom*) or MPPs (*Cd48*)(Figure S1B). Starting on day 14, but mostly at day 27, we found a continuum of differentiating cell states that we annotated using marker genes to seven major cluster groupings: granulocyte monocyte progenitors (GMP), megakaryocyte progenitors (Mk), Erythrocyte progenitors (Ery), Basophil progenitors (Ba), Monocyte progenitors (Mono) and Neutrophil progenitors (Neu)(Figure 1B,C and S1B - markers detailed on Supplementary Table 2). Even at day 27, we could still annotate thousands of cells as HSCs, confirming their expansion within these cultures (Figure 1C). This result suggested progressive differentiation and self-renewal from the initial pool of stem cells as previously reported (Wilkinson et al. 2019; Che et al. 2022).

Next, we leveraged LARRY barcoding to assess the clonal dynamics and cell fate choices of expanded HSCs over time. We found that expansion cultures gradually lose their clonality, despite initiating from highly pure EPCR+ HSCs, in line with recent reports (Figure 1D)(Zhang et al. 2024). Sister-cell splitting across independent wells confirmed the preferential expansion of specific clones, which correlated across wells higher than expected based on a null distribution obtained from a sampling simulation (Figure 1D). To describe the mechanisms leading to clonal selection, we visualized all clones detected in both D7 and D14 timepoints using “clone x state” heatmaps, which are colored based on the state-bias in each individual clone (Figure 1E). We observed a striking heterogeneity in lineage fate biases across clones even at D27 (Figure 1F, Figure S1C). More surprisingly, fate-biased HSCs showed the same fate-associated gene expression programs, which were similar to those previously identified in transplant or developmentally-traced native hematopoiesis (Figure S1D-E) (Rodriguez-Fraticelli et al. 2020; L. Li et al. 2023). These results indicate the preservation of clonal HSC heterogeneity and their transcriptional programs *ex vivo*.

We next compared the HSC clones that persist and expand until D27 (termed “high-fitness clones”) with the clones that are detected early but do not expand enough to be detected at later time points (termed “low-fitness clones”). Analysis of their fate properties indicated that outcompeted low-fitness clones had more rapid contribution to mature states (at D14) yet similar absolute numbers of HSCs, pointing towards intrinsic qualities (and not quantities) as the cause of fitness differences (Figure 1E, Figure S1F). To assess the early transcriptional differences associated with these differences in fitness, we used longitudinal retrospective state-fate analysis, comparing the HSC cell states at day 7 based on their fitness differences at day 27 (Figure 1G-H and Supplementary Table 3). High-fitness HSCs exhibited enriched expression of markers associated with self-renewal (*Procr*) and HSC-identity (*Mecom*, *Ly6a*, *Hlf*), as well as non-conventional retinoic signaling (*Rarb*), extracellular-matrix (*Sdc4*, *Mmp16*), synapses (*Dlg2*, *Ncam2*) and actin cytoskeleton regulation (*Fmnl2*, *Gimp*, *Palld*)(Figure 1H). High-fitness HSC clones also expressed higher levels of low-output and Mk-biased HSC signatures, as well as Skeletal morphogenesis signatures (e.g. *Tcf15*, *Myof*) (Figure 1I and and Supplementary Table 4). To validate these findings during expansion, we used the Tcf15-Venus mouse model, which enriches highly self-renewing HSCs (Rodriguez-Fraticelli et al. 2020). We sorted 500 CD45.1 wild-type E-SLAM HSCs and co-cultured them with either Tcf15^high^ or Tcf15^low^ E-SLAM HSCs (from a CD45.2 background). Tcf15^high^ HSCs consistently expanded whereas Tcf15^low^ cells were relatively outcompeted after 20 days in expansion cultures (Figure 1J). Taken together, these analyses indicate that *ex vivo* expansion cultures display a broad range of stable and dynamic fate behaviors, including intrinsic differences in fitness, which allows us to study how different stem cell clones respond upon acquisition of initiating cancer mutations.

### Sister-cell analysis and state-fate landscapes for Dnmt3a-R878H mutagenesis

To investigate how different cancer driver mutations influence stem cell fates, we developed a second set of lentiviral libraries constitutively expressing Cre-recombinase and a fluorescent reporter or the mock fluorescent reporter alone (Cre-P2A-mScarlet and mScarlet) and combined them with a conditional mouse model carrying a Cre-dependent Dnmt3a-R878H mutation, which is the mouse homolog of Dnmt3a-R882H, one of the most frequent driver mutations in acute myeloid leukemias (Loberg et al. 2019; Guryanova et al. 2016). We isolated HSCs from male and female mice, transduced them with differently-indexed T-Sapphire LARRY libraries, and then, on day 7, we profiled a part of the cells and split the remainder into a Cre or a mock labeling reaction, and then these were further split into separate wells that continued expansion independently(Figure 2A). The system allowed state-fate analysis for both wild-type and mutant clones arising from sister HSCs, which we termed scTRACK (simultaneous cell tracking of recombinase activation and clonal kinetics).

**Figure 2.**
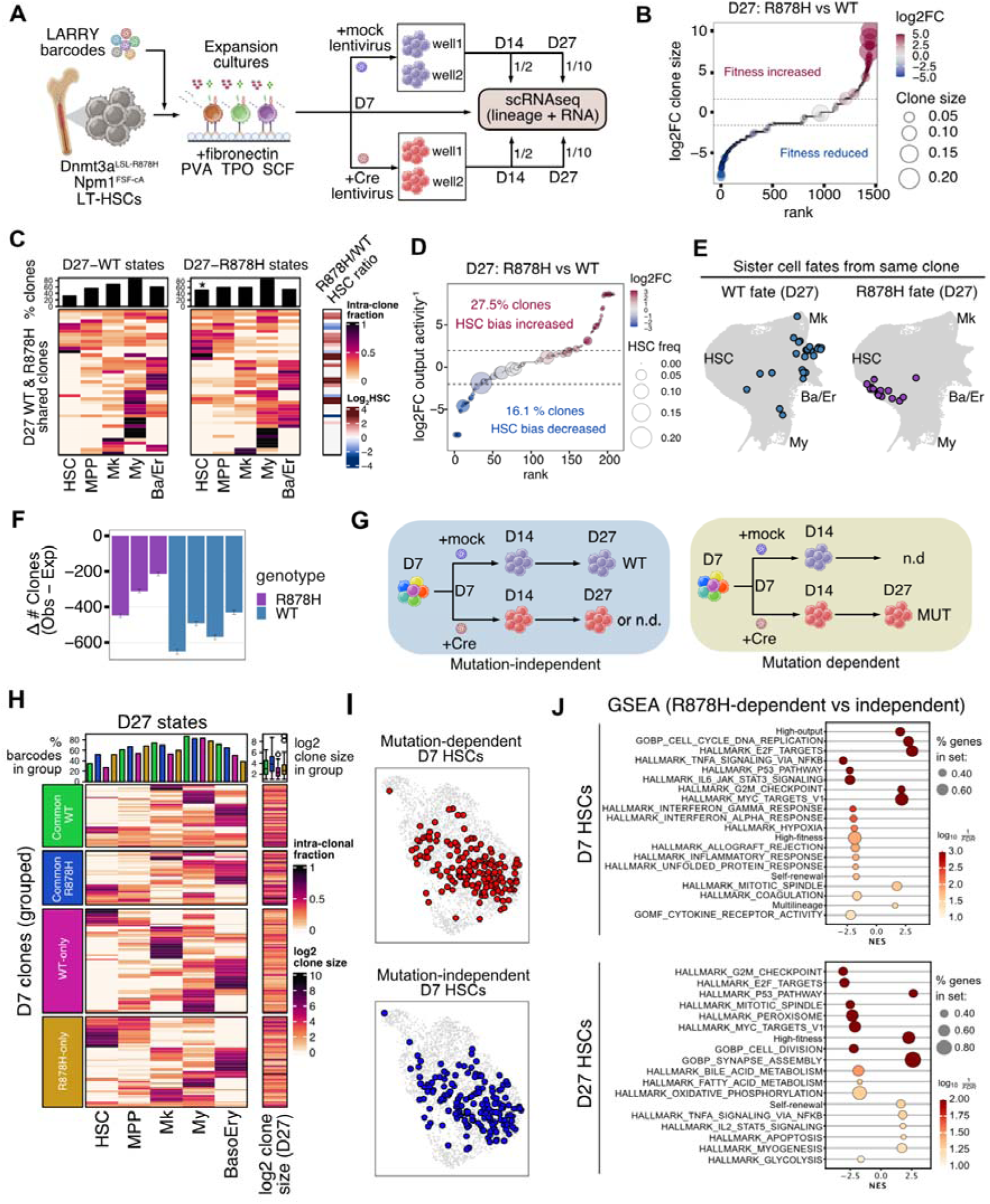
Sister-cell state-fate landscape of Dnmt3a-R878H mutagenesis. A) Experimental design for sister cell state-fate analysis in R878H mutagenesis. B) Waterfall plot showing log2 fold-change in clone size proportion for the same set of clones with and without the R878H mutation. C) State-bias heatmap of clones observed in both wt and R878H cultures at day 27. Log2 fold-change in HSC bias is shown on the right. Every row corresponds (in both heatmaps) to a single HSC clone. D) Waterfall plot showing log2 fold-change in HSC bias (inverse of output activity) for the same set of clones comparing R878H-mutant versus WT. E) UMAP of sister cell wild-type and mutant fates at day 27. The example shows a high-output clone that displays extreme low-output behavior with the R878H mutation. F) Difference between observed and expected number of clones at day 27. G) Schematic showing grouping of clones as mutation-dependent and mutation-independent. H) State-bias heatmaps of clones at day 27 separated into 4 clone-groups, depending on whether they are detected in both WT or R878H cultures (top heatmaps), WT-only or R878-only (bottom heatmaps). Clone size proportion is indicated in the right column. The top bar-plot indicates the proportion of clones with barcodes detected in each state (one column for each clone-group). I) UMAP of subsetted HSCs at day 7. Clones detected at day 27 are highlighted (mutation-dependent, top; mutation-independent, bottom). J) GSEA of hallmarks and HSC signatures for the comparison of R878H-dependent versus R878H-independent clones (at day 7, top; at day 27, bottom)

Similar to previous reports, we observed increased expansion of Dnmt3a R878H mutant cells (from here on R878H cells) in competitive cultures (Figure S2A), but did not identify major differences in their states (at the population level) compared to wild-type (WT) controls from the same mouse line (Figure S2B). After performing clonal analysis, we observed that HSC clones expanding significantly more with R878H also tended to have the largest clone sizes (Figure 2B). We then compared the clones that could be detected in both day 7 as well as day 27 in both WT and R878H cells by plotting their behaviors using state-bias heatmaps (Figure 2C). Sister WT/R878H clones displayed remarkably similar behaviors, even with a mutation and 20 days after splitting. Still, we noticed that most clones gained relatively more HSCs upon R878H mutation in comparison with the WT (Figure 2D), resulting in a net drop in the clonal output activity (Figure S2C). In some rare but notable cases, high-output multilineage clones even completely lost their output activity in the presence of the R878H mutation (Figure 2E), indicating that high-fitness stem-cells can be reprogrammed by the Dnmt3a mutation to gain self-renewal at the expense of differentiation. Thus, while these clones possess an inherent capacity for expansion irrespective of mutational status, the mutation drives further stemness capacity.

Considering these effects, we wondered whether the mutation could be altering the fitness capacity of intrinsically low-fitness clones. Comparing the clonality of R878H cultures with sampling simulations confirmed that R878H cultures maintained a relatively more polyclonal pool, suggesting that additional clones, normally outcompeted in the WT setting, persist only upon activation of the R878H mutation (Figure 2F). We next compared the clones detected only in R878H cells (mutation-dependent) with those detected only in WT or in both WT and R878H conditions (mutation-independent) (Figure 2G). We plotted state-bias heatmaps for day 27 states for each clone, split into groups based on their detection in WT and/or R878H cultures (Figure 2H). Notably, mutation-dependent R878H clones at day 27 behaved similarly compared to mutation-independent R878H clones, with increased HSC bias and a relatively larger size compared to WT-only clones. We next used retrospective state-fate analysis and compared the transcriptomes of day 7 HSCs based on their R878H mutation dependency at day 27 (Figure 2I). While we could not identify unique markers, gene set enrichment analysis (GSEA) of R878H-dependent versus independent clones indicated negative enrichment of self-renewal and high-fitness signatures, suggesting their origin in low-fitness HSCs (Figure 2J, top panel). Interestingly, post-mutation R878H-dependent HSCs displayed positive enrichment of high-fitness and self-renewal signatures (compared with mutation-independent clones), indicating the potent stemness reprogramming capacity of these driver mutations (Figure 2J, bottom panel) We further confirmed this by quantifying the single-cell fitness scores of R878H-dependent and independent HSCs at day 27 (Figure S2D). In addition to changes in HSC output bias, differential expression analysis and GSEA revealed that R878H HSCs (and MPPs) displayed reduced expression of early response genes, suggesting dampened inflammatory responses as an additional mechanism for their competitive expansion, in line with recent studies in clonal hematopoiesis (Figure S2E-F)(Serine Avagyan and Zon 2023; Jakobsen et al. 2023; S. Avagyan et al. 2021). Together, these results highlight the differential effect of this cancer driver mutation across different stem cell clones. While the R878H mutation can mildly enhance the stemness and expansion properties of high-fitness stem cell clones, it can reprogram the fates and states of low fitness stem cell clones, allowing them to survive and expand in *ex vivo* expansion cultures.

### The Flt3-Cre model recapitulates Dnmt3a-R878H mutagenesis in low-fitness HSCs in vivo

Next, we sought to validate the role of the Dnmt3a R878H mutation in low-fitness HSCs using a complementary approach. We took advantage of a modified version of the Flt3-switch approach, which normally labels developmentally restricted HSCs (Beaudin et al. 2016; Stonehouse et al. 2024). We speculated that we could identify low-fitness adult HSCs by combining the Flt3-Cre allele with the LSL-TdTomato reporter (tdTom), which is easier to recombine compared to mTmG or LSL-EYFP. We characterized the model at steady state in young mice using single-cell RNAseq profiling of tdTom+ and tdTom- LSK (Lin–c-Kit+Sca-1+) and LKs (Lin–c-Kit+ Sca1-)(Figure 3A,B). We verified that tdTom+ cells populated most clusters, while tdTom- were mostly restricted to HSC and MkP clusters, including bridging cells that suggest a direct Mk-restricted pathway (Figure 3C and S3A-C)(Haas et al. 2015; Yamamoto et al. 2018; Meng et al. 2023; Carrelha et al. 2018; Rodriguez-Fraticelli et al. 2018; Morcos et al. 2022). Approximately ∼50% of E-SLAM HSCs were labeled with tdTom+, suggesting that the Flt3-Cre tdTom model allows to separate the Mk-restricted and non-Mk-restricted hierarchies (Figure 3D). We next performed differential gene expression analysis on tdTom+ and tdTom- HSCs (Figure 3E). We found that tdTom+ cells showed relative downregulation of stem cell-associated genes like *Mecom* and positive enrichment for low-fitness and high output signatures, as well as IFNa signaling, and IL6/JAK/STAT3 signaling (Figure 3F). Thus, Flt3-Cre tdTom+ HSCs transcriptionally resemble low-fitness and high-output HSCs.

**Figure 3.**
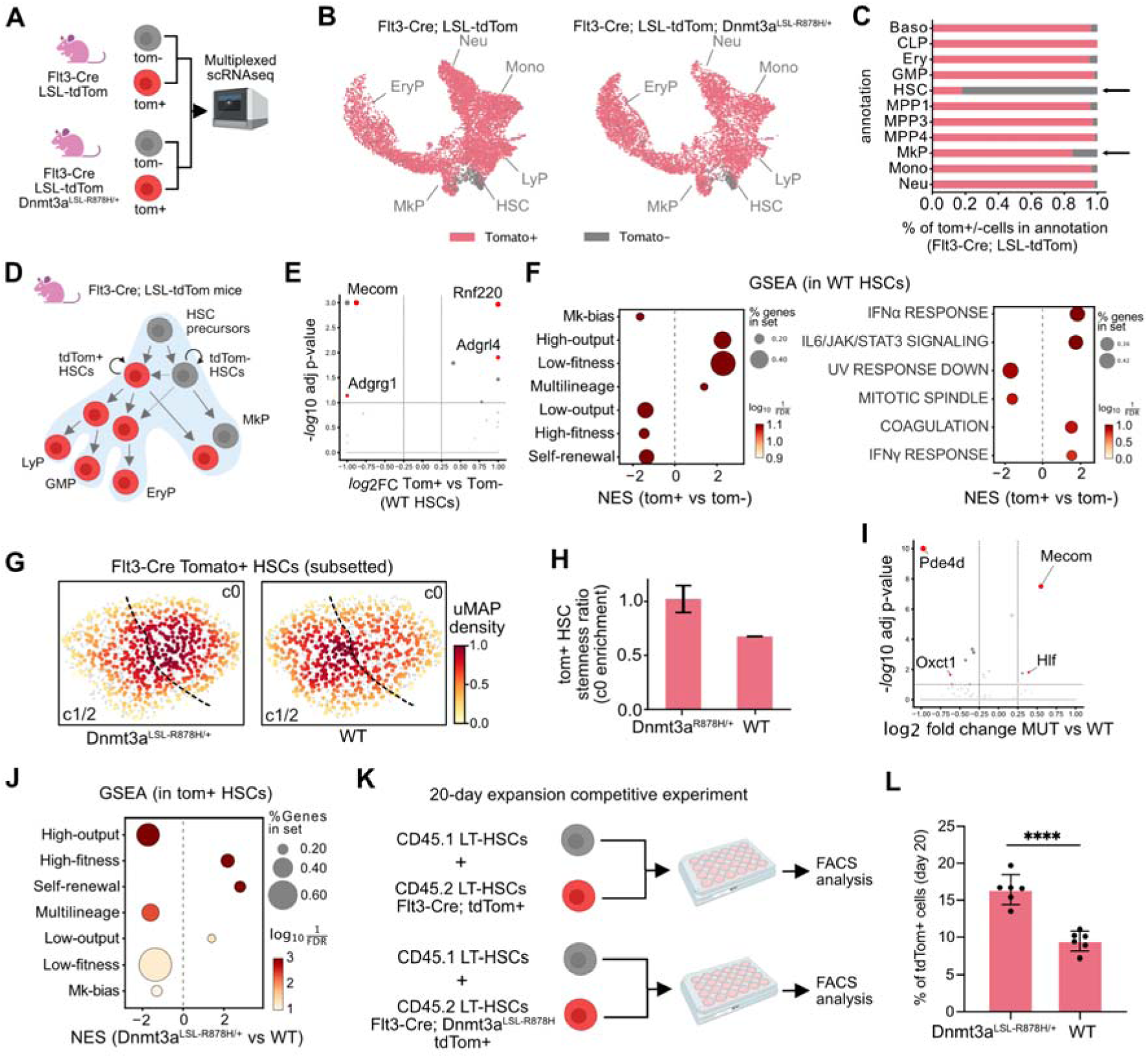
Dnmt3a-R878H rewires stem cell fitness in the Flt3-Cre HSC model. A) Experimental design for analysis of Flt3-Cre HSC models with 10x scRNAseq 3’. B) UMAP showing Flt3-Cre wt and Flt3-Cre R878H hematopoietic landscapes, with cells colored by TdTom expression. Annotated cluster groups are indicated. C) Barplot of an exemplary sample showing cluster distribution of TdTom+ and TdTom- cells. D) Scheme showing interpretation of Flt3-Cre model results. E) Volcano plot showing differential expression analysis of Flt3-Cre tdTom+ versus tdTom- HSCs. F) GSEA showing hallmarks and HSC signatures for Flt3-Cre tdTom+ versus tdTom- HSCs. G) UMAP of subsetted HSCs showing the distribution density. The dotted line marks the border between cluster 0 (higher stemness markers) and clusters 1 and 2 (higher cell-cycle and inflammatory response). H) Barplot showing relative enrichment in cluster 0 versus clusters 1 and 2. I) Volcano plot showing differential expression analysis of Flt3-Cre tdTom+ R878H-mutant versus Flt3-Cre tdTom+ wild-type HSCs. J) GSEA showing hallmarks and HSC signatures for Flt3-Cre tdTom+ R878H-mutant versus Flt3-Cre tdTom+ wild-type HSCs. K) Experimental design for CD45.1 competition experiment using *ex vivo* expansion cultures. L) Proportion of tdTom+ cells at day 20 comparing R878H versus wild-type.

To test the functional consequences of acquiring Dnmt3a R878H mutations in low-fitness HSCs, we generated Flt3-Cre; LSL-TdTomato; Dnmt3a-fl-R878H/+ mice and verified the specific mutagenesis in this tdTom+ HSC population by genotyping PCR (Figure S3D). Next, we compared the single-cell transcriptional landscape of R878H tdTom+ with wild-type tdTom+ hematopoiesis (Figure 3B). We observed a relative loss of myeloid cells and expansion of erythroid progenitors in the R878H, which is in line with results obtained in the whole-marrow Mx1-Cre R878H model, suggesting this phenotype can arise without mutagenesis in the most primitive tdTom- HSC compartment (Figure S3E). We then subsetted and re-clustered tdTom+ HSCs. R878H tdTom+ cells were relatively enriched in cluster c0, which expresses higher levels of stemness regulators and quiescence-associated genes (Figure 3G,H). In line with this, R878H tdTom+ HSCs showed upregulation of stemness genes compared to WT tdTom+ HSCs (*Hlf*, *Mecom*), and GSEA showed relative enrichment of low output, self-renewal and high-fitness signatures (Figure 3I,J). Finally, we mixed 500 Flt3-Cre tdTom+ HSCs (R878H or wt) with 500 (age-matched) CD45.1 wild-type HSCs using *ex vivo* expansion cultures (Figure 3K). We observed a ∼2-fold increased expansion of R878H tdTom+ HSCs over wild-type cells after 20 days, indicating the improved competitive advantage of R878H tdTom+ HSCs (Figure 3L). Altogether, these results validate our findings in *ex vivo* expansion cultures and demonstrate that the Dnmt3a-R878H mutation acquired *in vivo* within a low-fitness HSC population is sufficient to reprogram their transcriptional state and competitive behavior.

### Sister-cell analysis in Npm1c mutagenesis reveals clone-specific origins of mature versus primitive malignant states

Npm1c mutations are another frequent initiating alteration in AML and have been shown to endow stemness potential to mutant cells (Uckelmann et al. 2020; SanMiguel, Eudy, Loberg, Young, et al. 2022; Brunetti et al. 2018). To investigate the effects of Npm1c at clonal resolution, we developed a third set of lentiviral libraries constitutively expressing EGFP alongside Flpo recombinase or alone, and we then performed scTRACK using the same mouse model (Figure 4A). As with Dnmt3a, we again observed a very heterogeneous but significant expansion of Npm1cA mutant cells in competitive *ex vivo* expansion cultures (Figure 4B and S4A). This expansion was accompanied by the enrichment of Npm1c cells in an HSC-like state, which became conspicuous only at day 27, 20 days post-mutation (Figure 4C, S4B-C). Compared to WT cultures, Npm1c cultures maintained almost perfect clonality, losing less than 25% of the clones expected based on the stochastic sampling model (Figure 4D). Based on prior findings, we speculated that contribution from non-HSC clones (e.g. GMPs) could explain these results, but tracing back the sister-cell states of these clones at day 7 confirmed that most clones still originated in HSCs (Figure 4E). Based on our experience with Dnmt3a, we decided to classify HSCs as Npm1c-independent or -dependent and compared their origins at day 7. Npm1c-dependent HSCs showed a low fitness score at day 7, which became reversed at day 27, highlighting the potent fitness-programming effect of the Npm1c mutation (Figure S4D). Npm1c mutant cells showed expected gene expression changes compared to WT cells, including increased expression of HoxA cluster genes, proteasome and ribosomal components, and stemness markers (Figure S4E-I and Supplementary Table 3). To quantify clonal changes in fate behaviors in response to Npm1c-mutagenesis, we compared sister cell clones that had been profiled in both WT and Npm1c cultures at day 27 (Figure 4F-G). We noticed that Npm1c mutation tended to expand HSCs and reduce output activity and My bias, but this was highly variable across clones. Surprisingly, sister clones with low-output and high HSC content in the WT setting displayed more mature states and decreased the proportion of HSC-like cells after acquiring the Npm1c mutation. Conversely, sister clones with high-output properties in the WT showed the most primitive and differentiation-blocked behaviors in the context of the Npm1c mutation (Figure 4H and S4J). This response was highly heritable, with independently-mutated sisters of the original WT clone displaying similar behavior (Fig S4K). We next compared the gene expression profiles of clones that became more mature with Npm1c mutation (“HSC-decreased”) with clones that became more primitive (“HSC-increased”) (Figure 4I and Supplementary Table 3). HSC-decreased Npm1c clones expressed higher levels of mature malignant cell markers, including GMP genes (*Plac8*, *Mpo*) as well as various genes involved in AML function (*Zeb2*, *Plzf*)(H. Li et al. 2017; Ono et al. 2013). Intriguingly, mature-like malignant clones maintained expression of markers of their clone-of-origin, such as *Itsn1* or *Mir99ahg*, both of which are low-output/high-fitness HSC-markers. We complemented this analysis by comparing Npm1c clones based on the fitness score of their HSC of origin (at day 7), which revealed similar results (Figure S4L-M). Finally, we evaluated these clone-specific Npm1c signatures in human AML samples that were previously classified as “mature” or “primitive” (Mer et al. 2021; Naldini et al. 2023). Across two different cohorts, bulk or scRNAseq, we observed that “HSC-decreased” Npm1c clonal signatures were recapitulated in mature AML patients, suggesting an HSC-biased clonal origin in these leukemias (Figure 4J and S4N). To summarize, Npm1c mutations in low-output HSCs result in more mature malignancies, whereas mutations in high-output HSCs result in more primitive and aggressive expansions of malignant cells, contrary to our expectations. Together, our results point to a unique clone-of-origin mechanism for leukemia phenotypic heterogeneity, with pre-existing HSC states acting as a non-genetic substrate for the emergent properties of the malignant disease.

**Figure 4.**
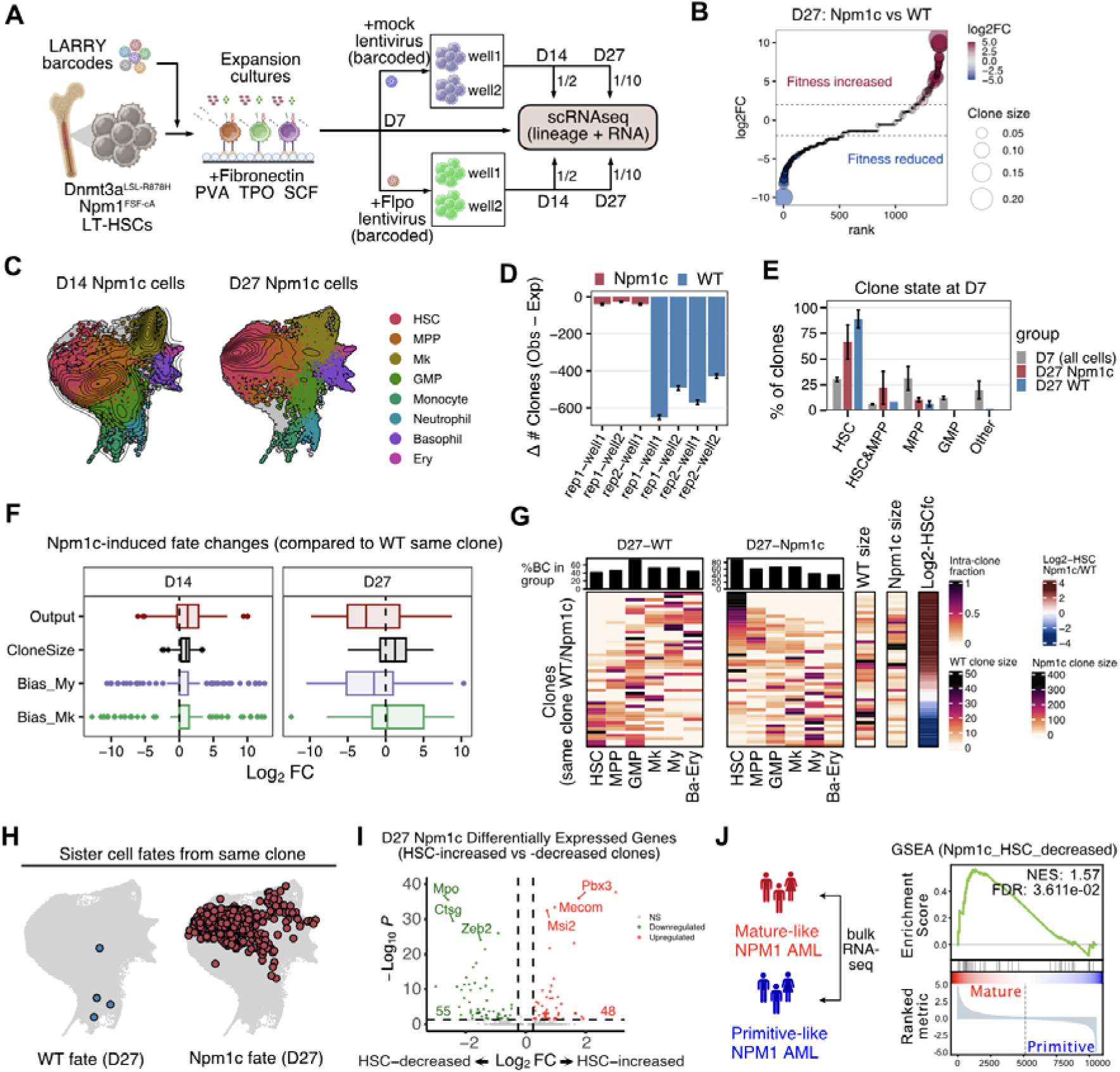
Pre-existing stem cell states-fates determine unique properties in malignant Npm1c clones. A) Experimental design for sister cell state-fate analysis in Npm1c mutagenesis. B) Waterfall plot showing log2 fold-change in clonal proportion for the same set of clones with and without the Npm1c mutation. C) Integrated UMAP showing Npm1c mutant cells (all clones combined) at day 14 (7 days after mutagenesis) and day 27 (20 days after mutagenesis), colored by cluster groups. D) Difference between observed and expected number of clones at day 27. E) Barplot showing distribution of day 7 states for clones observed at day 27 (shown separately for wt or Npm1c). The distribution of all day 7 states is shown for comparison. F) Boxplot showing sister-cell clonal behavior changes at day 27. Data is expressed as log2 fold change (Npm1c vs wt) in the indicated measurement (for all clones observed at day 27 in both Npm1c and wt cultures). G) State-bias heatmap of clones observed in both wt and Npm1c cultures at day 27. Clone size in each culture as well as log2 fold-change in HSC bias is shown on the right. H) UMAP of sister cell wild-type and mutant fates at day 27. The example shows a high-output clone that displays primitive leukemic behavior with the Npm1c mutation. I) Volcano plot results of differential gene expression analysis comparing Npm1c clones with HSC-increased versus Npm1c clones with HSC-decreased. Example markers are highlighted. J) GSEA of the Npm1c “HSC-decreased” signature (based on markers) performed on the list of mature vs. primitive NPM1 AML samples based on clustering of AML bulk-RNAseq data (Mer et al. 2021)

## Discussion

Cancer heterogeneity presents a significant challenge to oncologists. Phenotypic heterogeneity (plasticity, memory and noise) impacts critical therapeutic aspects of cancer biology, such as treatment resistance and clonal dominance. Recently, researchers have shown that phenotypic heterogeneity could be traced back to non-genetic variation at the level of individual cancer cells (Fennell et al. 2021; Goyal et al. 2023). The findings we present here further suggest that cancer heterogeneity may be partly influenced by subtle differences in the stem cell state of origin, leading to distinct responses even upon acquisition of identical cancer driver mutations. Classic cell-of-origin studies were limited by pre-existing tools and knowledge of cell type specific markers, which others and we have revealed to be insufficient to dissect the underlying heterogeneity of hematopoietic cell-types (Rodriguez-Fraticelli et al. 2020; Weinreb et al. 2020; Tian et al. 2021). Our systematic approach, combining sister-cell clone splitting with precise activation of cancer driver alleles, empowers a conceptual and methodological shift from the traditional cell-of-origin model to a more nuanced clone-of-origin paradigm, which embodies both lineage and state information.

At the population level, we observed that the Dnmt3a R878H mutation (R882H in humans) appeared to enhance the fitness and expansion of HSCs, consistent with prior studies (Loberg et al. 2019; Guryanova et al. 2016). Compared to their wild-type counterparts, R878H HSCs also exhibited reduced inflammatory signaling, which might add to their competitive edge, as recent CH studies in zebrafish, mice and humans have suggested (S. Avagyan et al. 2021; Serine Avagyan and Zon 2023; Schwartz et al. 2024; Jakobsen et al. 2023). However, at the individual clone level, R878H mutation had a more pronounced effect on high-output/low-fitness HSCs, which typically differentiate quickly and are outcompeted in expansion cultures, but, upon R878H mutation, gain enhanced stemness and fitness. Similarly, the Npm1c mutation consistently activated Pbx/Hoxa cluster expression across all clones, in line with its well described role in activating these developmental genes (Uckelmann et al. 2020; Brunetti et al. 2018; SanMiguel, Eudy, Loberg, Miles, et al. 2022). Yet, the individual clonal responses to this mutation varied greatly; unexpectedly, mutant sister cells from low-output HSCs tended to exhibit a more mature-like hierarchy. Previous studies in MLL-rearranged AML models have suggested that cancer cells inherit pre-existing functionalities of the cell of origin (George et al. 2016; Stavropoulou et al. 2016). However, our Npm1c results completely upset the notion that these state-biases should be maintained post mutation.

Our research also highlights the dynamic nature of these phenotypic transformations, which became apparent only 20 days post-mutagenesis, suggesting a role for cell divisions or secreted factors in culture media. Past work has also elucidated the role of genetic predispositions (Weinstock et al. 2023; Bick et al. 2020) and extrinsic regulation in the expansion of initiating (pre)-malignant clones (SanMiguel, Eudy, Loberg, Young, et al. 2022; Hormaechea-Agulla et al. 2021; Schwartz et al. 2024). In combination with our results, we would like to propose here the concept of a ‘clonal reaction norm,’ where clonal lineage, genetic background, driver mutations, and environmental factors collectively determine the fate and properties of a resulting cancer clone. Notably, our different Npm1c clones exhibited characteristics of various human NPM1 disease subtypes, indicating that understanding these norms for different cancer driver genes may have an impact on diagnostics and personalized treatments.

Finally, our experimental design should also be amenable to combinatorial and sequential mutagenesis. Future studies will focus on analyzing additional driver genes and systems beyond hematopoiesis. Looking ahead, we anticipate the development of new mouse models that can reproducibly generate tumors from clone-specific origins, offering a more accurate representation of the non-genetic tumor diversity that characterizes human patients to advance precision medicine efforts.

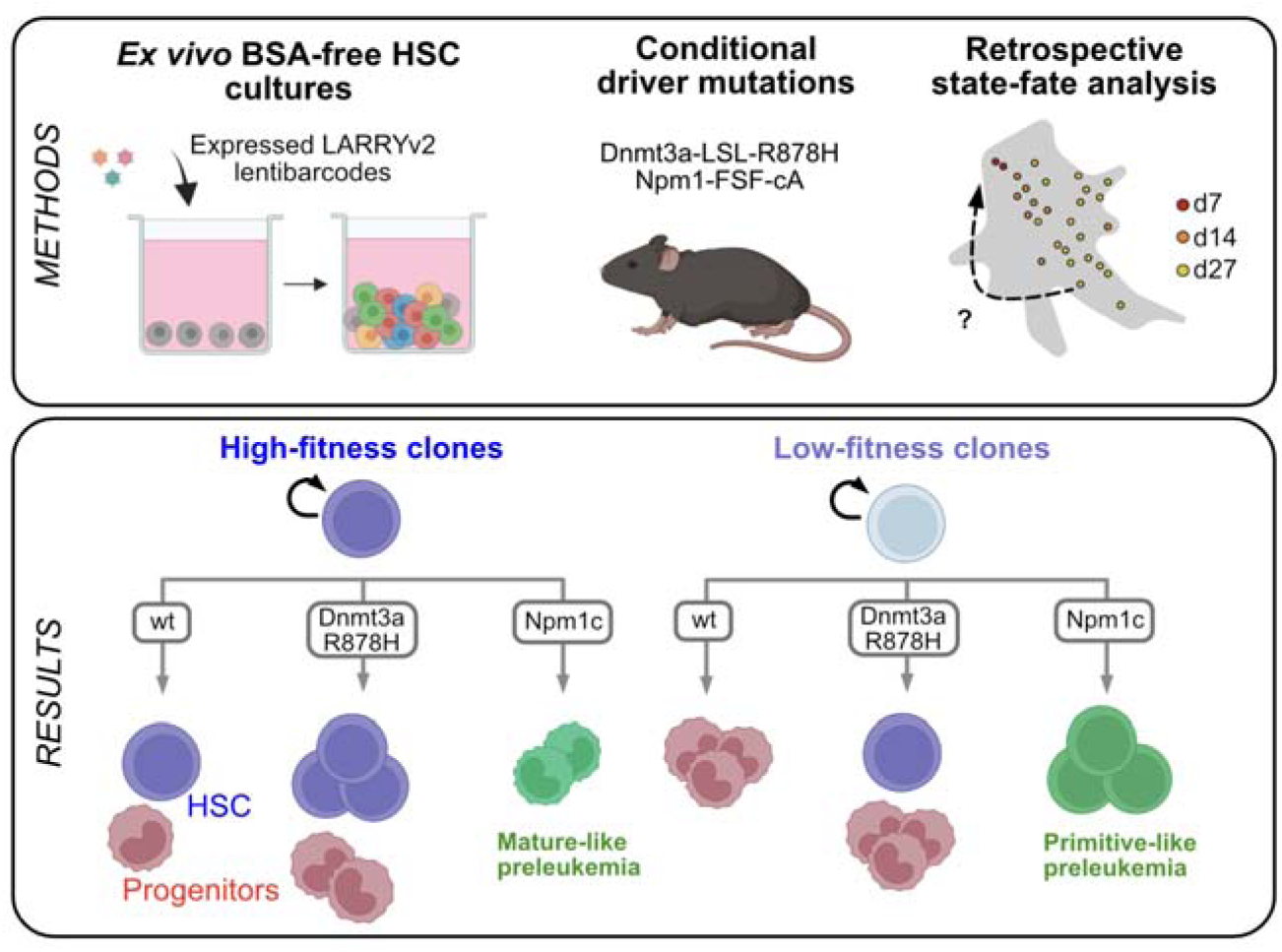

### Study limitations

These studies have been performed using mice, due to the accessibility of precision mutagenesis that can be achieved using genetically-engineered Cre/Flp-conditional mouse alleles. In the future, it is possible that Prime Editing or similar CRISPR-based precision editing technologies achieve the efficiencies to enable similar approaches in primary human HSCs (Geurts et al. 2021).

Our state-fate analysis uses *ex vivo* expansion cultures, to maximize our capacity to maintain and track many stem cell clones longitudinally, in separate environments, with and without mutations. We initially attempted to use stem cell transplantation to track cancer and wild-type clones, but these assays were underpowered, tracking too few clones, which is possibly due to mutant cell competition or niche heterogeneity. Our defined primary stem cell culture system is powerful as a proof of principle, but future technological implementations should address the role of niches and extrinsic components, which may take advantage of organoids or co-culture systems (Sommerkamp et al. 2021; Frenz-Wiessner et al. 2024; Khan et al. 2023).

Finally, our state-fate analysis relies on sister-cell splitting and clonal analysis, with sister states/fates serving as a proxy. This is particularly necessary due to the fact that our power relies on measuring 3 entities (mutant fate, wild-type fate and state) across thousands of cells. Novel technologies (such as Live-seq) are emerging to allow same-cell tracing without destruction, and may eventually have the capacity to perform STRACK-type studies at scale (Chen et al. 2022).

## SUPPLEMENTARY FIGURES

**Supplementary Figure 1.**
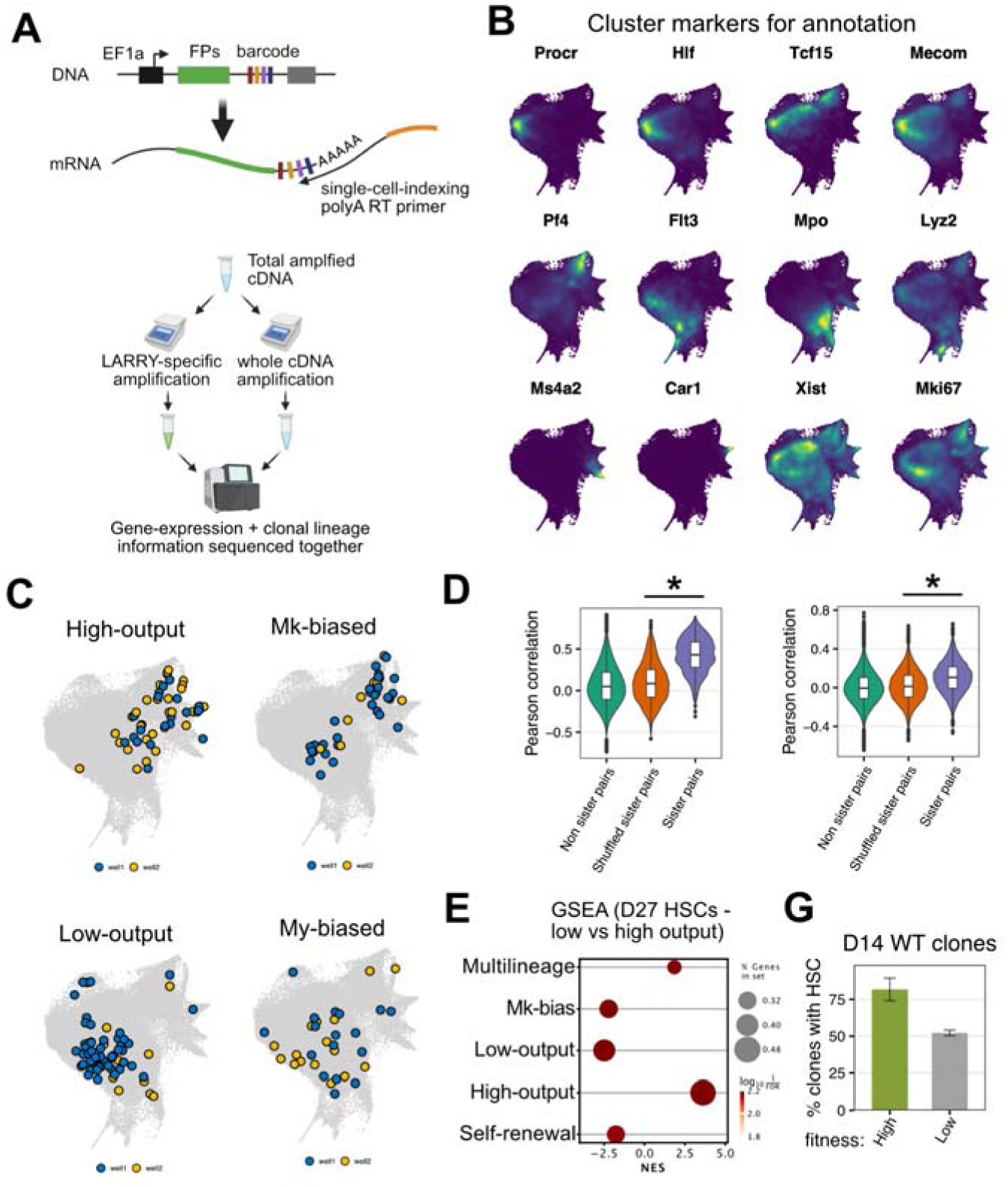
LARRY barcoding highlights a variety of clonal HSC behaviors in HSC expansion cultures. A) Principle for LARRY barcoding and state-fate analysis. Expressed barcodes can be captured efficiently with scRNAseq profiling methods. Then, single-cell indexed cDNA libraries are split in two parts: one for amplifying the whole transcriptome (GEX library) and one for amplifying the clonal barcode specifically (LARRY library). These libraries are sequenced together and then demultiplexed to feed to the CloneRanger pipeline. B) UMAP showing scaled (min-to-max) expression for various cluster markers. C) Integrated UMAP showing similar clonal behaviors in WT clones split across both wells (well 1 - blue; well 2 - yellow). D) Pearson correlation comparing sister cell pairs (cells from the same clone) versus shuffled or non-sister pairs at day 7 and day 27. * p<0.001 (permutation test) E) GSEA of HSC signatures in D27 high-output HSCs versus low-output HSCs. F) Percentage of clones with an HSC at D14 (comparing high versus low fitness)

**Supplementary Figure 2.**
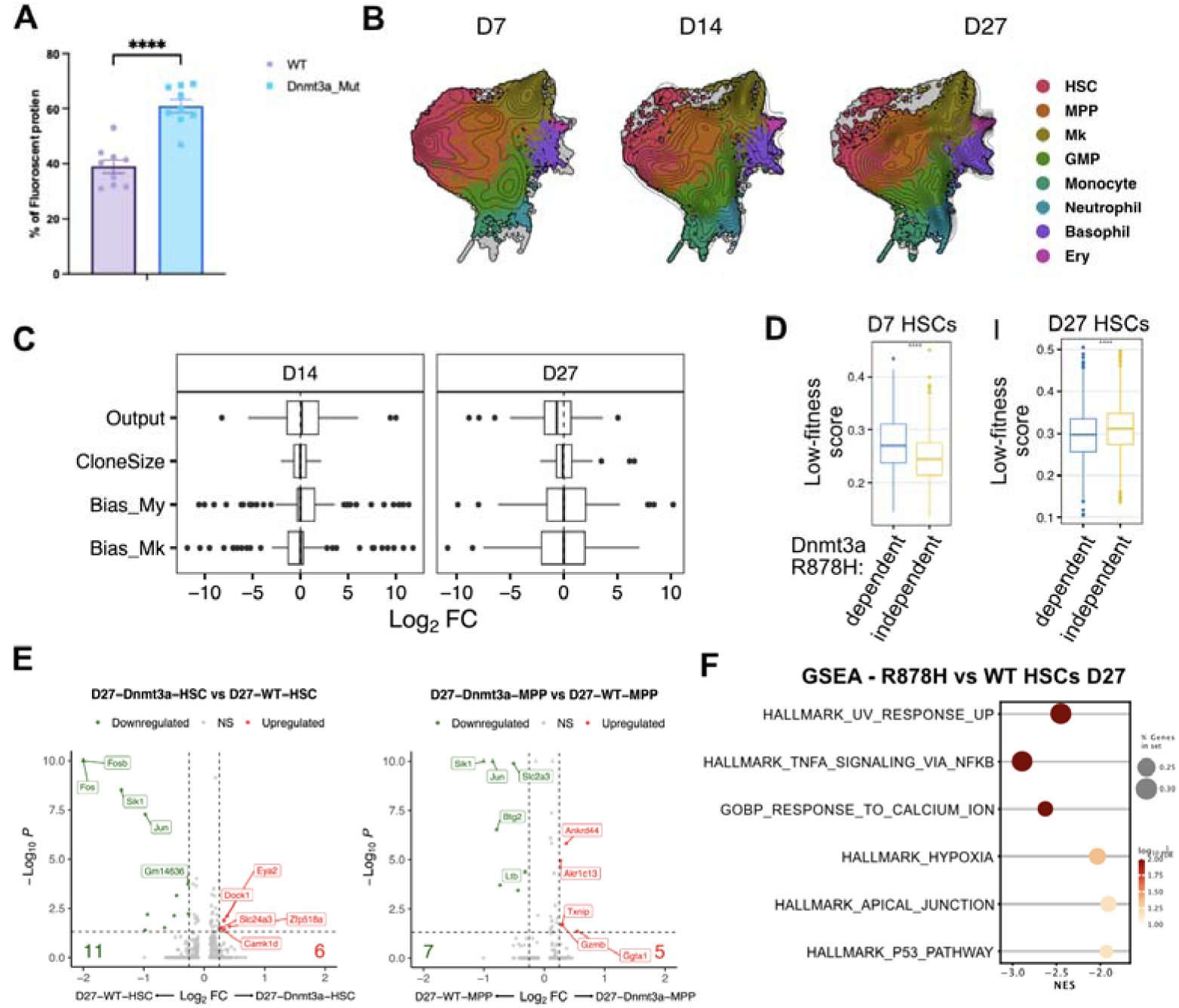
Differences in R878H effects at the population or clonal level. A) Percentage of Dnmt3a-R878H mutant or wild-type cells after 20 day *ex vivo* expansion culture. HSCs were isolated from Dnmt3a-LSL-R878H mice and transduced with mock or Cre lentivectors. Then, 500 Cre mutant cells (or 500 mock cells) were co-cultured with 500 CD45.1 cells. Cre or wt mutant cells were measured at day 20.**** p<0.001 (two-sample t-test, n=9) B) UMAP showing cluster annotations for Dnmt3a-R878H cells at different timepoints. C) Boxplots showing changes in clonal behaviors (R878H versus WT) for the same clones measured in both cultures at day 14 and day 27. D) Boxplots showing low-fitness HSC signature score differences across R878H-dependent and independent HSCs (at day 7 and day 27) ***p<0.001 (wilcoxon test) E) Volcano plot showing differential expressed genes comparing R878H vs WT HSCs or MPPs (at day 27). F) GSEA of significant MsigDB hallmarks and GO terms in D27 R878H v WT HSCs.

**Supplementary Figure 3.**
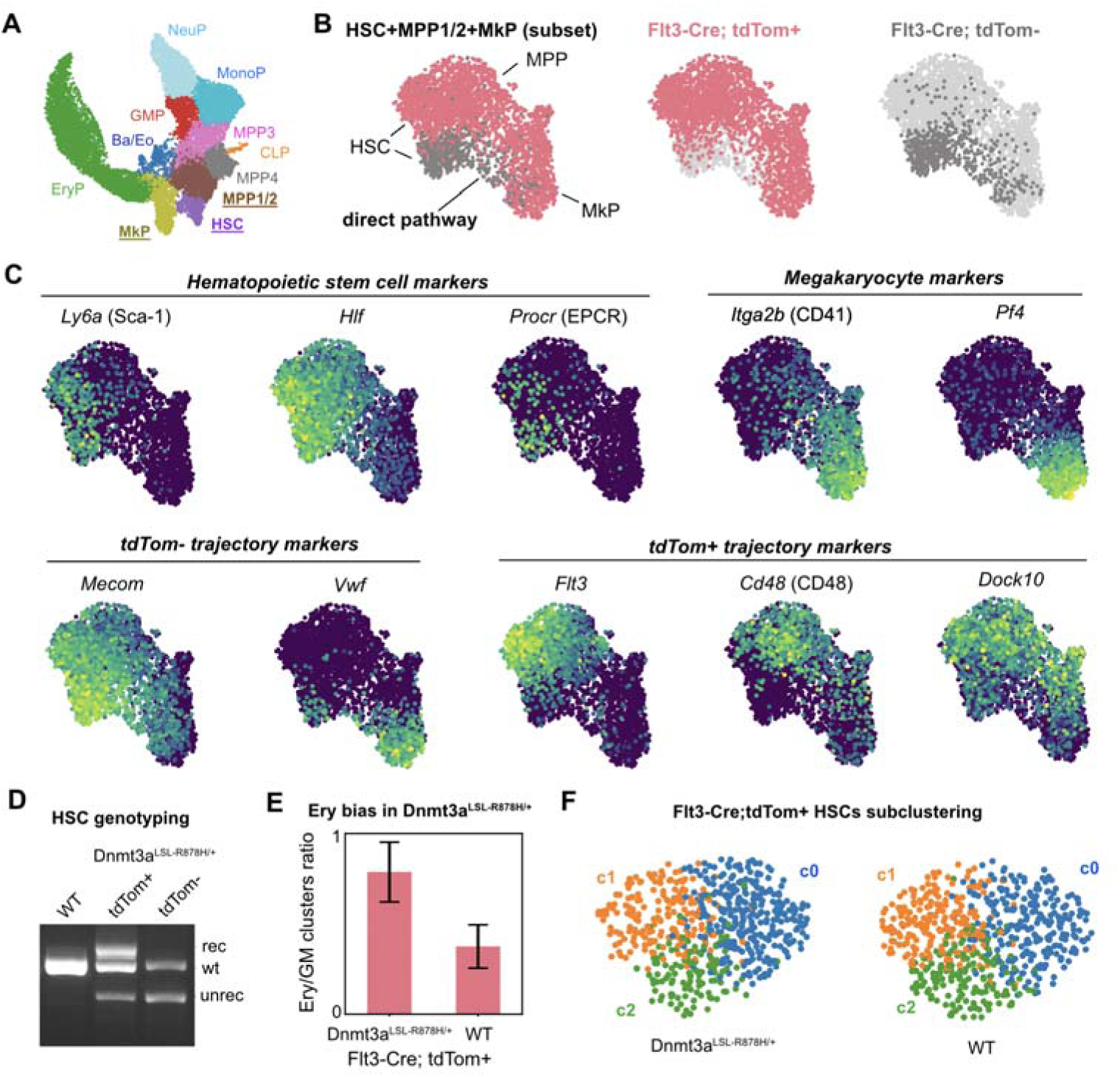
Flt3-Cre model labels different stem cell compartments and trajectories. A) UMAP showing HSPC cluster groupings as profiled from Flt3-Cre LSL-tdTom mice. B) UMAP of subsetted HSC, MPP and MkP clusters, showing tdTom+ (red) and tdTom- (gray) cells. A trajectory of tdTom- cells suggests evidence for negative labeling of a direct Mk pathway from the most primitive HSCs. C) UMAP of subsetted HSC/MPP/MkPs showing scaled expression of different HSC, MPP and MkP markers. D) Genotyping of R878H mutation in Flt3-Cre Dnmt3a-LSL-R878H LSL-tdTom model. Cells were sorted and genotyped as in Loberg et al. E) Erythroid bias in Dnmt3a-LSL-R878H mutant tdTom+ cells, compared to WT. Plot shows ratio between the proportions of the Ery to GM clusters in each replicate. F) Subclustering of subsetted tdTom+ HSCs showing 3 distinct clusters. Cluster 0 shows enrichment in stemness markers (e.g. *Mecom*).

**Supplementary Figure 4.**
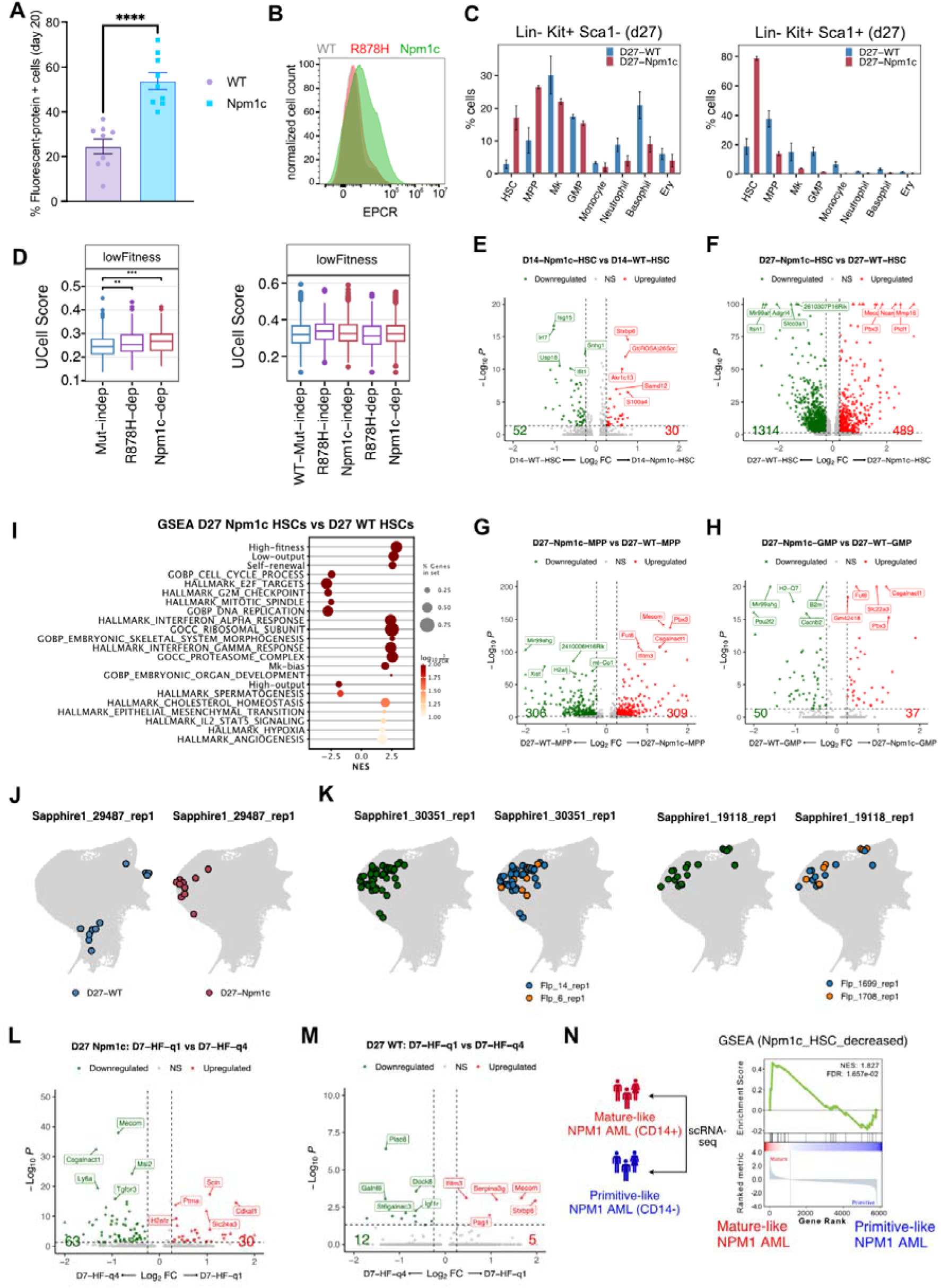
Additional analyses on Npm1c malignant clones. A) Percentage of Npm1c mutant or wild-type cells after 20 day *ex vivo* expansion culture. HSCs were isolated from Npm1-FSF-cA mice and transduced with mock or Flpo lentivectors. Then, 500 Flpo mutant cells (or 500 mock cells) were co-cultured with 500 CD45.1 competitor cells. Flpo or wt mutant cells were measured at day 20 by FACS.**** p<0.001 (two-sample t-test, n=9). B) Example histogram of EPCR expression at day 27 in WT, R878H and Npm1c cultures. C) Annotation of LSK and LK states in Npm1c versus wild-type clones at day 27. D) Volcano plot showing differences between Npm1c and wild-type HSCs at day 14. E) Volcano plot showing differences between Npm1c and wild-type HSCs at day 27. F) Volcano plot showing differences between Npm1c and wild-type MPPs at day 27. G) Volcano plot showing differences between Npm1c and wild-type GMPs at day 27. H) GSEA of significant HSC signatures, MsigDB hallmarks and GO terms in D27 Npm1c v WT HSCs. I) Ucell scores for the Low-fitness signature at day 7 and day 27 for WT-only (Mutation-independent), R878H or Npm1c HSCs (Mutation-dependent). J) UMAP of sister cell wild-type and mutant fates at day 27. This additional example shows a high-output clone that displays a more primitive leukemic behavior with the Npm1c mutation. K) UMAP of sister cell mutant fates (2 different mutant subclones of the same pre-mutation WT clone) at day 27. This example shows low-output primitive leukemic behavior for both subclones detected. L) Volcano plot showing differences between Npm1c clones derived from high-fitness (q1 - quartile 1) versus low-fitness (q4 - quartile 4) HSCs at day 7. Notice enrichment of stemness genes in q4-derived clones. M) Volcano plot showing differences between WT clones derived from high-fitness (q1 - quartile 1) versus low-fitness (q4 - quartile 4) HSCs at day 7. Notice enrichment of stemness genes in q1-derived clones. N) GSEA of the Npm1c “HSC-decreased” signature performed on mature vs. primitive NPM1 AML samples as profiled based on AML scRNAseq data (Naldini et al. 2023). Samples were assigned to each group based on CD96 expression (primitive marker) and CD14 expression (mature marker).

## ACKNOWLEDGEMENTS

I.S. was supported through the European Union’s Horizon 2020 research and innovation program under the Marie Skłodowska-Curie grant agreement No 945352.

D.F.P was supported through the Marie Skłodowska-Curie grant agreement No 101109276.

A.R.F is supported by the Cris Foundation Excellence Award (PR_EX_2020-24), the ERC Starting Grant MemOriStem (101042992), the Spanish National Research Agency (PID2020-114638RA-I00), the Agencia de Gestio d’Ajuts Universitaris i de Recerca (AGAUR, 2017 SGR 1322), and the CERCA Program/Generalitat de Catalunya. A.R.F. acknowledges support from the Institut Catalá de Recerca i Estudis Avançats (ICREA), the Ministry of Science Ramon y Cajal Fellowship, and the LaCaixa Junior Fellows Incoming Fellowship.

The authors would like to acknowledge the technical assistance of David Fernández, José Ignacio Pons, and Freddy Monteiro from the Functional Genomics Core Facility at IRB Barcelona (single-cell library preparations and sequencing). We also acknowledge assistance from the Flow Cytometry and Cell Sorting Core Facility at the University of Barcelona (CCIT-UB) and from the facilities from the Parc Cientific de Barcelona (PCB). The authors also wish to thank mentors and colleagues for various helpful discussions. Illustrations were created with Biorender.

## Author Contributions

I.S. performed molecular biology, library preparations, cell culture, and animal experiments. D.F.P. generated the analytic pipelines and performed bioinformatic, single-cell sequencing, and statistical analyses. A.R.F and P.S. assisted in the generation of the bioinformatic pipeline and analysis. A.R.F., I.S., and D.F.P conceptualized the project design. A.R.F. designed the LARRY libraries. I.S. generated all the vectors and produced the lentiviral libraries with assistance from lab members. I.S. and D.F.P. wrote the manuscript with help from A.R.F. All authors provided feedback and input to finalize the manuscript text.

## RESOURCE AVAILABILITY

### Lead contact

Further information and requests for resources and reagents should be directed to and will be fulfilled by the lead contact, Alejo E. Rodriguez-Fraticelli

### Materials availability

Plasmids generated by this study are available upon request and will be deposited to Addgene and the European Plasmid Repository.

### Data and code availability

Code and data objects are available at figshare (DOI: 10.6084/m9.figshare.25822948). Raw and processed single-cell RNAseq, LARRY barcoding and TotalSeq™ data generated for this study will be released at Gene Expression Omnibus under the accession number GSE266232.

## MATERIALS AND METHODS

### Mice and Animal Guidelines

All procedures involving animals adhered to the pertinent regulations and guidelines. Approval and oversight for all protocols and strains of mice were granted by the Institutional Review Board and the Institutional Animal Care and Use Committee at Parque Científico de Barcelona under protocol CEEA-PCB-22-001-ARF. The study follows all relevant ethical regulations. Mice were kept under specific pathogen-free conditions for all experiments.

### Hematopoietic stem cell isolation

Following euthanasia, bone marrow was harvested from the femur, tibia, pelvis, and sternum through mechanical crushing, ensuring the retrieval of most HSCs. The collected bone marrow cells were then sieved through a 100-μm strainer and cleansed with a cold ‘Easy Sep’ buffer containing PBS with 2% fetal bovine serum (FBS), followed by lysis of red blood cells using RBC lysis buffer (Biolegend, Catalog no. 420302). At first, mature lineage cells were selectively depleted through the Lineage Cell Depletion Kit, mouse (Miltenyi Biotec, Catalog no. 130-110-470), while the resulting Lin- (lineage-negative) fraction was then enriched for c-Kit expression using CD117 MicroBeads (Miltenyi Biotec, Catalog no: 130-091-224). These cKit-enriched cells were washed, blocked with FcX and stained with following fluorescently labeled antibodies: APC anti-mouse CD117, clone ACK2 (Biolegend catalog no. 105812), PE/Cy7 anti-mouse Ly6a (Sca-1) (Biolegend, catalog no. 108114); Pacific Blue anti-mouse Lineage Cocktail (Biolegend, catalog no. 133310); PE anti-mouse CD201 (EPCR) (Biolegend, catalog no. 141504); PE/Cy5 anti-mouse CD150 (SLAM) (Biolegend, catalog no. 115912); APC/Cyanine7 anti-mouse CD48 (Biolegend, catalog no. 103432).

### HSC *ex-vivo* expansion cultures

*Ex-vivo* cultures of HSCs were done under self-renewing F12-PVA-based conditions as described previously (Wilkinson et al. 2019). To this end, cell-culture activated 96-well flat-bottom plates were coated with a layer of 100 ng/ml fibronectin (Bovine Fibronectin Protein, CF Catalog: 1030-FN) for 30 minutes at room temperature. Following the sorting process, HSCs were transferred into 200 µl of complete HSC media supplemented with 100ng/ml recombinant mouse TPO and 10ng/ml recombinant mouse SCF (PeproTech Recombinant Murine TPO Catalogue Number: 315-14; PeproTech Recombinant Murine SCF, Catalogue Number: 250-03) and grown at 37°C with 5% CO2. During lentiviral library transduction, the first media change took place 24 hours post-transduction. All other protocol steps followed the guidelines provided in (Wilkinson et al. 2020).

### Construction of lentiviral pLARRY vectors

The construction of barcoded libraries was executed by a previously established protocol (https://www.protocols.io/view/barcode-plasmid-library-cloning-4hggt3w). First, the T-Sapphire, Scarlett, or EGFP coding sequences, and the EF1a promoter sequences were PCR amplified from pEB1-T-Sapphire, pmScarlet_NES_C1, and pLARRY-EGFP with primers homologous to the vector insertion site in a custom synthetic lentiviral plasmid backbone (Vectorbuilder, Inc) using Gibson assembly (Gibson Assembly® Master Mix, NEB, Ref. E2611L). For recombinase lentivirus libraries, iCre or Flpo recombinase was PCR amplified together with EGFP and Scarlett with primers homologous to the vector insertion site in a custom synthetic lentiviral plasmid backbone and cloned using Gibson assembly. After magnetic-bead purification, ligated vectors were then transformed into NEB10-beta electroporation ultracompetent E.coli cells (NEB® 10-beta Electrocompetent E. coli, NEB, Ref.C3020K) and grown overnight on LB plates supplemented with 50 μg/mL Carbenicillin (Carbenicillin disodium salt, Thermo Scientific Chemicals Ref. 11568616). Colonies were scrapped using LB medium and pelleted by centrifugation. Plasmid maxipreps were performed using the Endotoxin-Free Plasmid Maxi Kit (Macheray Nagel), following the manufacturer’s protocol. pEB1-T-Sapphire was a gift from Philippe Cluzel (Addgene plasmid 103977). pLARRY-EGFP was a gift from Fernando Camargo (Addgene plasmid 140025). pmScarlet_NES_C1 was a gift from Dorus Gadella. Additional reagent details are in Supplementary Table 5.

### Barcode lentivirus library generation and diversity estimation

To barcode pLARRYv2 plasmids and generate a library, first a spacer sequence flanked by EcoRV restriction sites was cloned into the plasmid after the WPRE element of the vector. Custom PAGE-purified single-strand oligonucleotides with a pattern of 20 random bases and surrounded by 25 nucleotides homologous to the vector insertion site were synthesized by IDT DNA Technologies (Supplementary Table 5). The assembly of these components and subsequent purification steps were carried out through Gibson assembly (Gibson Assembly® Master Mix, NEB, Ref. E2611L). Six electroporations of the bead-purified ligations were performed into NEB10-beta E.coli cells (NEB® 10-beta Electrocompetent E. coli, New England BiolabsEB, Ref.C3020K) utilizing a Gene Pulser electroporator (Biorad). Subsequently, after transformation, the cells were incubated at 37 degrees for 1 hour at 220 rpm. Post-incubation, the transformed cells were plated in six large LB-ampicillin agar plates overnight at 30°C. Colonies from all six plates were collected by scraping with LB-ampicillin and then grown for an additional 2h at 225 rpm and 30 °C. Cultures were pelleted by centrifugation, and plasmids were isolated using the Endotoxin-Free Plasmid Maxi Kit (Macheray-Nagel), following the manufacturer’s protocol. For estimating diversity, barcode amplicon libraries were prepared by PCR amplification of the lentiviral library maxiprep using flanking oligonucleotides carrying TruSeq read1 and read2 adaptors using 10 ng of the library (Supplementary Table 5). We used the minimal number of cycles that we could detect by qPCR to avoid PCR amplification bias (10-12 cycles). After bead purification, 10 ng of the first PCR product was used as a template for a second PCR to add Illumina P5 and P7 adaptors and indexes (Supplementary Table 5). Two independent PCRs were sequenced on an Illumina NovaSeq 6000 S4 platform (Novogene UK) to confirm diversity after correction of errors through collapsing with a Hamming distance of 4. After collapsing, libraries were confirmed to contain at least 50 million different barcodes, with enough diversity for uniquely labeling up to 100,000 HSCs with a minimal false-positive rate. Lentivirus production and HSPC transduction were performed as described in (Weinreb et al. 2020).

### Single-cell encapsulation and library preparation for sequencing

For scRNA sequencing and subsequent plating, cells were pipetted up and down gently a few times to be dissociated into single cells and transferred to a 1.5 mL microtube. The well was then washed with prewarmed PBS to collect all the possible remaining cells. Cells were then concentrated by centrifugation at 800 g for 8 minutes. Washed cells were then blocked with FcX, and stained with the E-SLAM stem cell antibodies panel, to confirm expansion of E-SLAM cells. Live cells were then sorted based on fluorescent reporter expression. Part of the sample as specified in text was then taken for constructing a single-cell library using Chromium Single Cell 3’ Reagent Kits (v3) following the manufacturer’s guidelines (10X Genomics). The remaining part was then plated back for further expansion in culture. To minimize the impact of batch effects on sequencing, we multiplexed different conditions leveraging the unique barcode pattern of our libraries together with Biolegend TotalSeq™ anti-mouse hashing antibodies (Supplementary Table 5), enabling the simultaneous preparation of libraries representing all experimental conditions in a single reaction for each day of sampling.

Following the reverse transcription of mRNA and first-strand cDNA amplification, 100 ng of the cDNA libraries were used as templates to amplify LARRY barcodes by nested PCR similar to the protocol described in (Weinreb et al. 2020). The first PCR used forward primer (Pre-Enrichment forward) CTG AGC AAA GAC CCC AAC GAG AA together with the corresponding 10x Genomics dual index TruSeq reverse primer using the following programs 1, 98 C, 3 min; 2, 98 C, 20 s; 3, 58 C, 15 s; 4, 72 C, 20 s; 5, repeat steps 2–4 08 times; 6, 72 C, 3 min; 7, 4 C, hold. The PCR products were then purified with a 0.8:1 ratio of Ampure XP beads. Purified PCR products were then subjected to a second PCR using the forward primer (Trueseq_LARRY) GTG ACT GGA GTT CAG ACG TGT GCT CTT CCG ATC TGC TAG GAG AGA CCA TAT GGG ATC and the corresponding 10x dual index Truseq reserve primer, following program 1, 98 C, 3 min; 2, 98 C, 20 s; 3, 58 C, 15 s; 4, 72 C, 20 s; 5, repeat steps 2–4 08 times; 6, 72 C, 3 min; 7, 4 C, hold. The final PCR products were then purified by a 0.8:1 ratio of Ampure XP bead: PCR products, were indexed using the 10x dual index TruSeq kit, and sequenced using Illumina NovaSeq or NextSeq.

### scRNA-seq data processing and calling of lineage barcodes

Generation of single-cell matrices for gene expression and LARRY lineage barcodes was performed using *cloneRanger*, an in-house developed pipeline (https://github.com/dfernandezperez/cloneRanger) to process 10XGenomics single-cell RNA-seq together with LARRY barcoding. The pipeline is based on Snakemake(Köster and Rahmann 2012) and the use of Docker/singularity containers to allow for reproducibility and easy deployment of the code.

For each sample, fastq files from gene expression (GEX), LARRY and TotalSeq™ tags were processed using cellranger v7.0.0 with default parameters. However, since cellranger only collapses barcodes that are 1 hamming distance apart, prior to the execution of cellranger, fastq files containing LARRY barcodes were processed using UMICollapse(Liu 2019). This allowed us to collapse all barcodes which were 4 hamming distance units apart or less, similar to the procedure used by (Weinreb et al. 2020; Rodriguez-Fraticelli et al. 2020). In particular, the UMICollapse was executed with the following parameters: *“fastq -k 4 --tag”*. Finally, in order to run cellranger in feature barcode mode with LARRY and TotalSeq™ sequences, we created a reference library csv file by extracting all detected LARRY barcodes across all collapsed fastq files, together with TotalSeq™ sequences. A reference library file was created for each individual sample and given as input to cellranger, executed with default parameters. All the code and steps performed by the pipeline are available in the *cloneRanger* github page.

A Seurat(Hao et al. 2024) object containing single-cell count matrices from GEX, LARRY and TotalSeq™ counts was created with the function *Read10X* from Seurat. Finally, cell doublets were removed with scDblFinder(Germain et al. 2021) using default parameters and TotalSeq™ sequences were demultiplexed with the function *hashedDrops* from the DropletUtils R package(Lun et al. 2019) with default parameters.

The assignment of LARRY barcodes to individual cells was performed by *cloneRanger* similarly to (Weinreb et al. 2020; Rodriguez-Fraticelli et al. 2020): first, we generated a filtered LARRY matrix by removing barcode UMIs that were sustained by less than 5 sequencing reads (this information is stored in the *molecule_info.h5* file generated by cellranger). Then, we further filtered the LARRY matrix by removing all barcodes with less than 4 UMIs. After filtering, barcodes were assigned to individual cells as following: (a) for cells in which only one barcode was detected after filtering, that barcode was assigned, (b) for cells in which more than one barcode was detected post-filtering, the top barcode with higher UMI counts was assigned and (c) for cells in which there were ties in the top barcode, no barcode was assigned. Our barcode calling strategy was developed to minimize mixing cells from different clones at the expenses of having higher chances to split real clones into subclones.

### Single-cell integration, clustering and annotation

To integrate scRNA-seq samples, we applied the *IntegrateLayers* Seurat v5 workflow to test multiple integration algorithms (sample time point was used as batch variable): Harmony, Reciprocal PCA, Canonical Correlation Analysis and Joint PCA. After supervising the results from all algorithms, we decided to use Reciprocal PCA to integrate the single-cell GEX matrices. We followed a standard Seurat pipeline with some minor modifications. Raw counts were normalized with the function *NormalizeData* and the top 3000 variable genes were selected. From those, we removed all ribosomal and mitochondrial genes, as well as genes that correlated with cell cycle genes (*Ube2c*, *Hmgb2*, *Hmgn2*, *Tuba1b*, *Ccnb1*, *Tubb5*, *Top2a*, *Tubb4b*, pearson cor of 0.1 or more), as performed by (Weinreb et al. 2020). Then, filtered variable genes were used to compute the top 50 reciprocal-PCA components. The kNN graph was computed using the function *FindNeighbors* setting the number of neighbors to 20, which was extended to 30 for the generation of UMAP components. Clusters were generated with the function *FindClusters* with a resolution of 0.3 and the Louvain algorithm.

Annotation of cell types was performed using known gene markers from the literature. A summary of the main markers for every cluster are shown in Supplementary Figure 1B and the whole list of markers for every cluster, computed with *FindAllMarkers* from Seurat, are listed in Supplementary Table 2.

### Quantification and classification of HSC clonal behaviors

Clone x state heatmaps from Figures 1, 2, and 4 were generated as follows: for every clone, we computed the number of cells detected across every cluster (cell type), generating a clone-by-cluster matrix A. In A, each A_ij_ represents the number of cells from the cluster *j* detected in the clone *i*. After generating A, in order to account for cell type abundance heterogeneity, we column-normalized the matrix by the total number of cells from each cell type, generating a B matrix. Finally, to compare clonal fates between clones of different sizes, we row-normalized the B matrix to obtain, for every clone, the fraction of cells present in each cell type (intra-clone fraction score). HSC clonal behaviors were quantified as described in (Rodriguez-Fraticelli et al. 2020). [explain briefly]. To determine statistically significant biased clones (clones representing a higher proportion of a specific cluster than expected from random cell sampling) we applied a Fisher’s exact test as described in (Biddy et al. 2018) accounting for clone size.

### Sister cell similarity analysis

To calculate the sister cell similarity scores shown in Supplementary Figure 1D, we subsetted individually the cells corresponding to each timepoint (Day 7 and D27) and proceeded as follows: The top 2000 variable genes were selected and, as described above, ribosomal, mitochondrial and genes correlating with cell cycle genes were filtered out. These filtered variable genes were scaled and used as input to calculate the top 70 principal components (PCs). The cell-by-PC matrix (obtained with the function *Embeddings* from Seurat) was used as input for the R *cor* function selecting *pearson* as the correlation metric. This procedure generated a cell-by-cell similarity matrix that was split into: (a) all pairs of sister cells, (b) all pairs of non-sister cells, and (c) all pairs of sister cells in which the barcode label was previously shuffled. To assess the statistical significance between the average Perason coefficient of these 3 groups, a permutation test with 1000 simulations was performed. Briefly, the average correlation of each group was compared to a random distribution of sister similarity scores generated by shuffling the larry barcodes prior to the generation of the cell-by-cell similarity matrix across 1000 iterations.

### Stochastic sampling model of clonal selection

To calculate the expected clonality across our experimental time course, assuming that all clones have equal fitness, we developed a null clonal selection model based on sampling with a binomial distribution. This model recapitulates the different sampling events (sample splitting, well splitting, sampling of cells for scRNA-seq, fraction of cells encapsulated in library preparation) and measured cell expansion (from D7 to D14 and D14 to D27). It makes two key assumptions for simplicity: no cell death (based on our limited observation of apoptotic-like events during culture) and similar proliferation probabilities for all cell types. We applied sequential sampling calls with replacement (using the R function *sample(x, replace = TRUE*)) to model, from the initial distribution of clones sizes detected at D7, the following steps -using empirical data for each individual well replicate-: 1) fraction of cells lost in encapsulation for single-cell RNAseq at D7, 2) fraction of cells split into WT and mutant samples at D7, 3) transduction efficiency of secondary LARRY barcoding, 3) fraction of cells split into different wells, 4) cell expansion from D7 to D14, 5) fraction of cells used for D14 sequencing, 6) cell expansion from D14 to D27, 7) fraction of cells sampled for D27 sequencing and 8) fraction of cell recovery (encapsulation) from D27 library preparation. The output of the model is a list of the expected clones detected at D27 and their corresponding sizes. We ran 1000 iterations of the model, from which we calculated an average expected clone size. The distribution of clones sizes from the model was also used to calculate the expected clone size correlation between wells at D27.

### Single-cell differential gene expression and signature scores

All gene differential expression analyses, unless otherwise specified, were performed using MAST (Finak et al. 2015) within the function *findMarkers* from Seurat using the replicate information as the only latent variable. This was done in order to mask differences in gene expression between male and female cells, which corresponded to replicate 1 and replicate 2, respectively.

### Gene set enrichment analysis

For each comparison, we created pre-ranked lists based on log2 fold-change differences in gene expression obtained from scRNAseq or bulkRNAseq analysis. These pre-ranked lists were analyzed using *gseapy* version 1.1.1 (Fang, Liu, and Peltz 2023). For gene sets, we used signatures shown on Supplementary Table 4 (HSC signatures) or MsigDB gene sets (mouse hallmarks or gene-ontology terms). Data from AML patients were obtained from previous studies (Mer et al. 2021; Naldini et al. 2023). For the scRNAseq dataset, patients were categorized as “mature” or “primitive” based on the ratio of % cells expressing CD14 versus % cells expressing CD96. Patients with CD14/CD96 ratio >1 were labeled as “mature”, while CD14/CD96 ratio <1 were labeled as “primitive”. For the bulkRNAseq dataset, we used the differential expression list output in the available code (comparing the “mature” patient cluster with the “primitive” patient cluster).

### Statistical methods

Statistical analysis was performed using the tests as indicated throughout the text. Generally, Wilcoxon rank-sum tests were used for statistical significance except where indicated.

## SUPPLEMENTARY INFORMATION

**Supplementary Table 1. Summary statistics of clones.** This table contains the summary statistics for each library and replicate in the study, including number of clones, average size, number of cells and percentage of the culture sampled.

**Supplementary Table 2. Cluster markers and groupings.** This table contains the markers used to annotate the clusters and group them into annotations.

**Supplementary Table 3. Differential expressed gene analyses.** This table contains all the differential gene expression analysis results. Different comparisons are under different tabs.

**Supplementary Table 4. Clonal HSC signatures used.** This table contains the list of genes used for gene set enrichment analysis.

**Supplementary Table 5. Reagents tables.** This table contains the lists of reagents used, including genotyping primers and LARRY barcode sequences.

